# Metagenomics Analyses of Cellulose and Volatile Fatty Acids Metabolism by Microorganisms in the Cow Rumen

**DOI:** 10.1101/414961

**Authors:** Lijun Wang, Xin Jiang, Hongjian Xu, Yonggen Zhang

**Affiliations:** College of Animal Science and Technology, Northeast Agricultural University, Harbin, 150030, China

**Keywords:** forage-to-concentrate ratio, fiber decomposition, volatile fatty acid production, fibrolytic bacteria, gene expression

## Abstract

The purpose of this study was to evaluate the effects of different forage-to-concentrate (F:C) ratios (7:3 high-forage, 3:7 high-concentrate) on rumen microflora and fiber degradation mechanism. Compared with the high-concentrate (HC) group, the high-forage(HF) group showed improved fiber degradation and a sustained high level of carboxymethyl cellulose (CMCase), β-glucosidase and β-xylosidase activities, but the total VFAs decreased. Among bacteria at the family level, *Lachnospiraceae* and *Succinivibrionaceae* in HF groups were 2-fold and 4-fold more abundant than in the HC group, respectively. A KEGG analysis revealed that succinate-CoA synthetase (EC: 6.2.1.5) and propionate-CoA transferase (EC: 2.8.3.1) leading directly to propionate production were more abundant in HC group. Conversely, butyryl-CoA dehydrogenase (EC: 1.3.8.1) was directly related to butyrate production and was higher in the HF group. A gene expression analysis showed that the relative content of *Fibrobacter succinogenes* and *Butyrivibrio fibrisolvens* was higher in the HF group and contributed more to fiber degradation and VFA production. *Prevotella ruminicola, Selenomonas ruminantium*, and *Veillonella alkalescens* contributed more to starch degradation and propionate production, which relative content was higher in the HC group. This research gave a further explanation of the fiber degradation parameters and microbiota under different F:C ration. The fiber-degrading bacteria in the roughage group have a high content level, and the corresponding cellulase activity is also high. These results supported the potential of diets for microbial manipulation, which can increase feed digestibility and explored new fibrinolytic bacteria.

**IMPORTANCE:** The forage of the cow’s feed occupies a large proportion. The shortage of high-quality forage in cow breeding has become an important factor limiting the China’s dairy industry. The effective measure is to improve the utilization of low-quality forage. Based on traditional nutrient metabolism, the reasons for the effects of roughage on the growth and metabolism of dairy cows can be explored, but the metabolic mechanism is not well analyzed, and the further utilization of forage is also limited. Metagenomics has proven to be a powerful tool for studying rumen microbial structures and gene function. This experiment used metagenomics to study the metabolism of cellulose and volatile acids in the rumen. Our research showed that different forage-to-concentrate shifted the composition of microorganisms and the activity of enzymes, resulting in different metabolic pathways of volatile fatty acids. This work provides a background for microbial community composition and further use of forage.

## 1. INTRODUCTION

The rumen ecosystem is recognized as a natural bioreactor for highly efficient degradation of fibers, and rumen microbes have an important effect on fiber degradation (1). The rumen provides anaerobic conditions and redox potentials that favor microbial growth and expression of fiber-degrading enzymes, which allows rumen microorganisms to breakdown cellulosic plant materials to meet ruminants’ daily energy requirements by producing volatile fatty acids (VFAs). The degradation of carbohydrates in the rumen can be divided into two stages Fig.1 (1, 2). For example, cellulose is first hydrolyzed into cellobiose by ruminal microorganisms and then is further hydrolyzed to glucose, after which the glucose continues to be fermented to produce pyruvate (3-5). Finally, a series of fermentation reactions produce VFAs (6). These processes are all performed through the actions of rumen microbial enzymes. The changes of F:C ratios in the diet significantly affect the number and type of rumen microorganisms and then affect the end products of fermentation (7, 8). Previous studies show that the content of *Fibrobacter* and *Ruminococcus* (both are mainly Fibrinolytic bacteria) increased when dietary fiber increased in the rumen (9, 10). The type of diet directly causes changes in the rumen environment, causing changes in the rumen microbial population, population structure and enzyme activity. Mcallan et al. (1994) and Ribeiro et al.(2015) (11, 12) studied the effects of rumen fermentation on cows with high-forage and high-concentrate diets, with the results indicating that total VFA and NH_3_-N were not affected. When concentrates in the diet increased, the acetate and butyrate were significantly reduced, but the propionate significantly increased.

**Fig. 1.**
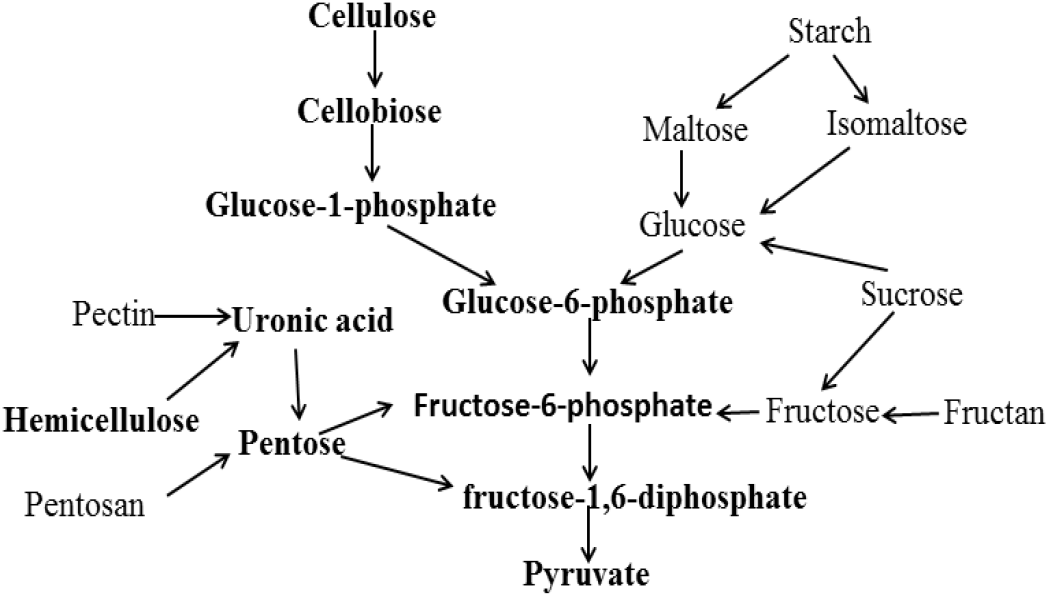
Degradation of polysaccharides to produce pyruvate (6)

Although much work has been done to investigate the effects of F:C in the diet of dairy animals on metabolism, most of the studies focused on the metabolism of nutrients or the production of volatile acids, which does not provide a systematic change, i.e., from cellulose fermentation to end-product VFAs and the variation of the microbial population during fermentation. The ruminal microbial ecosystem is an effective model for degrading fiber; therefore, it is important to understand how ecosystems develop and operate when shifting the diet of the host. Therefore, we combine metabolism and metagenomics to explore the changes from cellulose to end product VFAs and the changes of cellulose in rumen. At the same time, we also quantified major functional bacterial species using quantitative PCR to assess the role of these organisms in adapting to different diets (HC and HF).

However, we studied how rumen bacteria change and adapt to different rumen environments in different diets. Therefore, it is of practical significance to study the competition between fibrinolytic bacteria (the result of competition and its internal mechanism).

## 2. MATERIALS and METHODS

**2.1 Experimental Design, Animals and Sample Collection**

Twelve ruminally cannulated, lactating Holstein cows averaging 3.2 ± 0.70 (mean ± SE) years of age (range = 0.9 years) were used in this experiment. Cows were placed in individual tie stalls in a temperature-controlled room. Six animals (lactating cow) were fed a F:C ratio of 7:3, and the other six animals (dry cow) were fed a F:C ratio of 3:7. Diet composition and nutrients were shown in Supplement Tab. 1. Animals were fed once daily at 8:00 h and allowed ad libitum consumption at 110% of the expected intake for four weeks before being sampled.

**Table 1.**
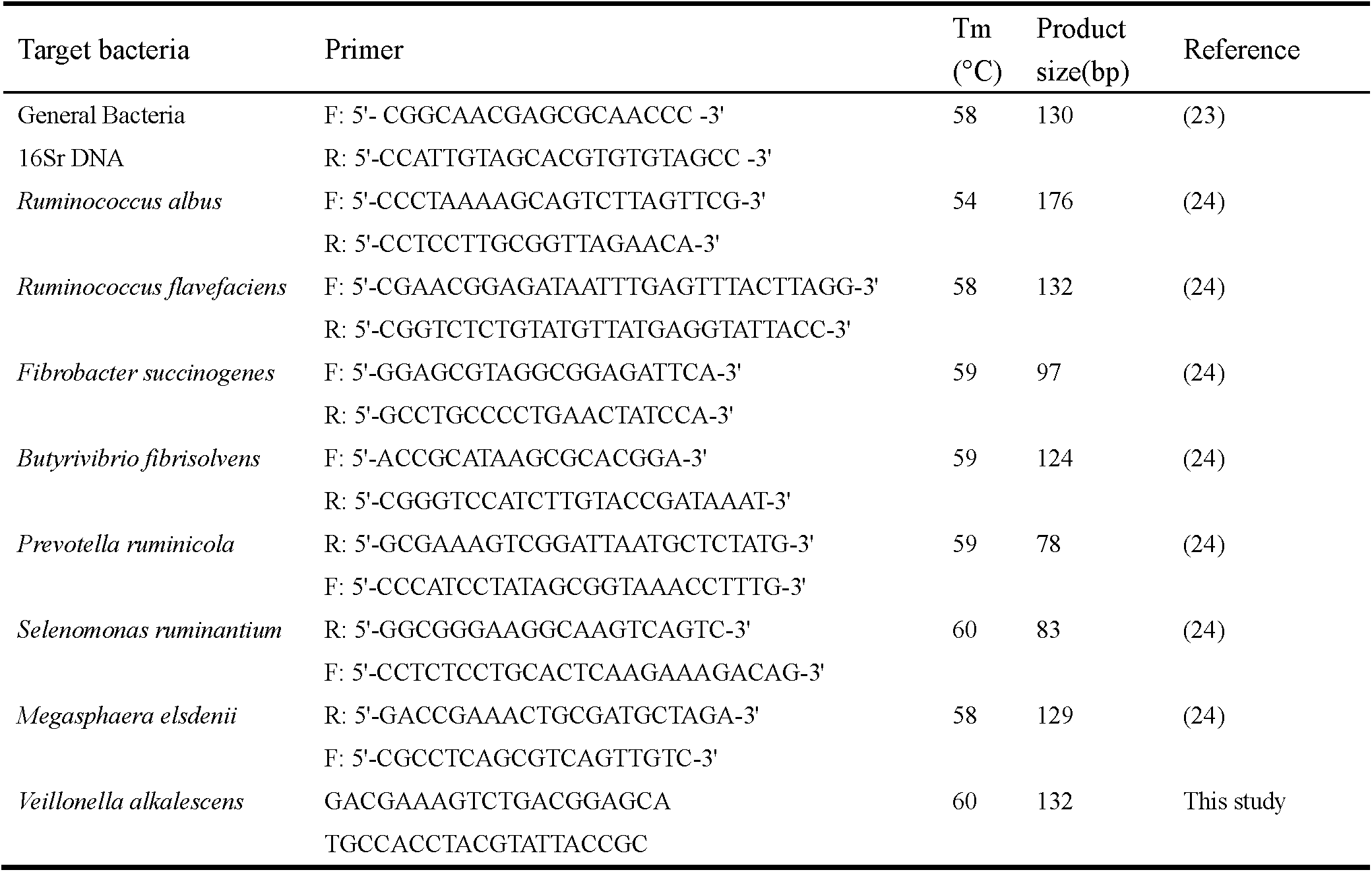
Primers used for real-time PCR quantification

**Table 2.** Fermentation products of carbohydrates in the rumen

|  | Con | Forage | SEM | P-value |
| --- | --- | --- | --- | --- |
| pH | 5.76 | 6.14 | 0.0719 | 0.0208 |
| NH<sub>3</sub>-N, mg/dL | 11.94 | 16.51 | 0.5363 | <.0001 |
| Acetate, mM | 86.41 | 80.09 | 2.1322 | 0.5929 |
| Propionate, mM | 41.82 | 23.52 | 1.0549 | <.0001 |
| Butyrate, mM | 16.89 | 12.96 | 0.4438 | 0.0002 |
| Valerate, mM | 2.77 | 0.67 | 0.0234 | <.0001 |
| Isobutyrate, mM | 1.24 | 4.17 | 0.1179 | <.0001 |
| Isovalerate, mM | 1.92 | 0.78 | 0.0537 | <.0001 |
| TVFA, mM | 151.04 | 130.18 | 3.5658 | 0.0033 |
| Acetate: Propionate | 2.07 | 3.76 | 0.0711 | <.0001 |
| Lactate (mmol/L) | 6.61 | 8.48 | 0.48 | 0.0213 |

<sup>a</sup>TVFA: Total volatile fatty acid

**Table 3.** The chemical composition of diets and rumen contents

|  | Con | Forage | SEM | P-value |
| --- | --- | --- | --- | --- |
| Ruminal contents |  |  |  |  |
| Cellulose | 22.07 | 23.13 | 2.14 | 0.8013 |
| Hemicellulose | 22.83 | 31.88 | 7.22 | 0.5442 |
| Acid detergent lignin (ADL) | 12.24 | 17.93 | 1.96 | 0.1616 |
| Diets |  |  |  |  |
| Cellulose | 13.16 | 16.69 | 0.25 | <.0001 |
| Hemicellulose | 16.02 | 28.88 | 1.25 | <.0001 |
| Acid detergent lignin | 6.00 | 8.61 | 0.20 | <.0001 |

**Table 4.**
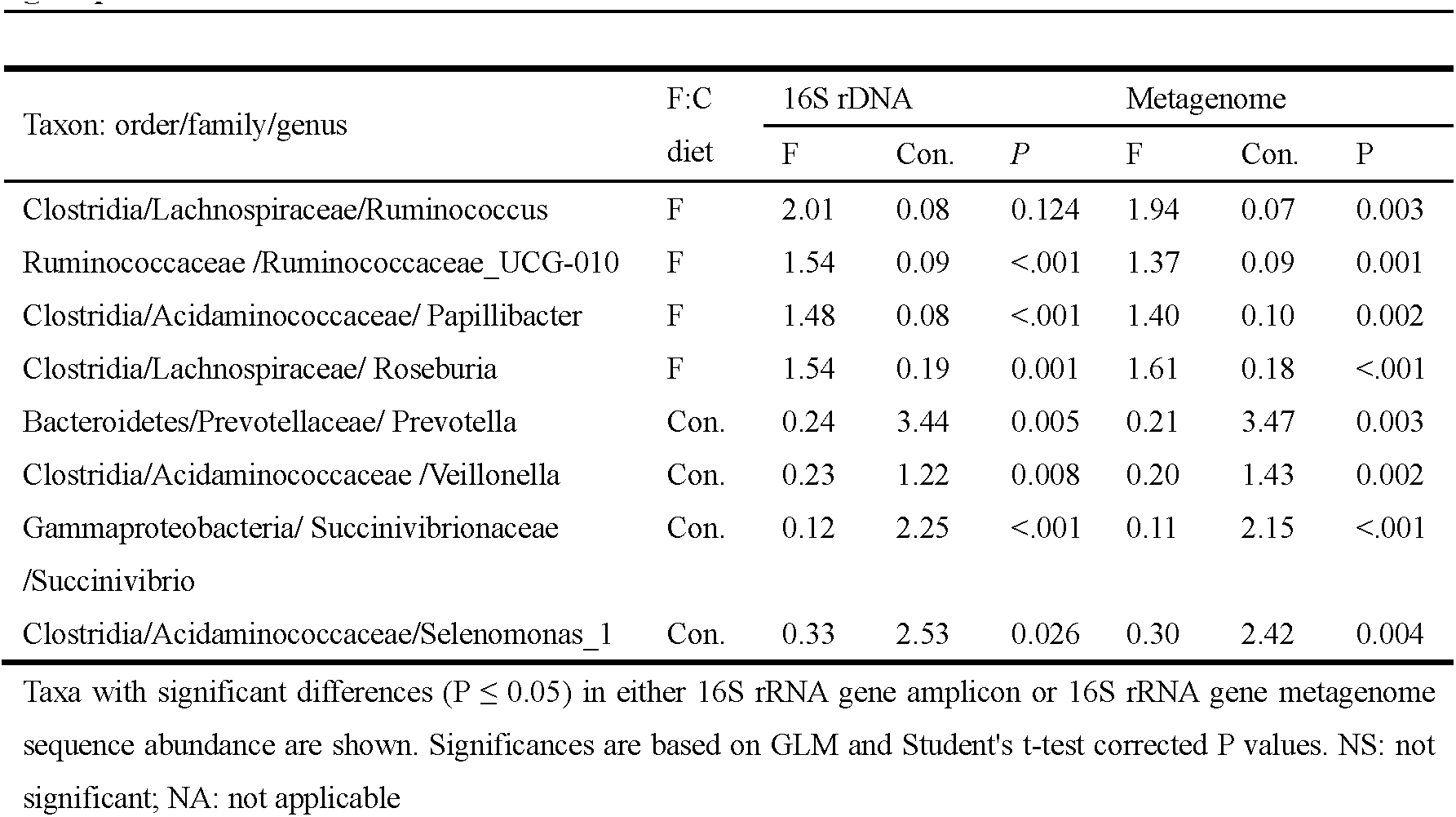
Bacterial taxa (97% sequence similarity) with taxonomy assigned to the highest possible resolution, differing in mean relative abundance (%) between the HF and HC groups measured.

### Compliance with Ethical Standards

**Conflicts of interest** All authors declare that they have no conflict of interest.

**Ethics approval** All animal studies were conducted according to the animal care and use guidelines of the Animal Care and Use Committee of Animal Science and Technology College, Northeast Agricultural University.

### 2.2 Sample Collection and Measurements

Rumen content samples were collected 4 h after feeding via a ruminal fistula. Representative rumen content samples were collected from each animal, and the solid from the liquid through four layers of cheesecloth were sampled. One part of each homogenized pellet was mixed with RNAlater (Ambion, Texas, USA), a reagent that protects and stabilizes cellular RNA. One part of each homogenized pellet with freshly prepared metaphosphoric acid (25% w/v; 1 mL) was added to 5 mL of filtered rumen fluid and vortexed, which was used to measured volatile fatty acids (VFAs). All samples were placed in liquid nitrogen within five min and were then taken to the laboratory and stored at −80°C until further testing.

For the determination of VFAs, samples with metaphosphoric acid were thawed at room temperature and then centrifuged (12,000 g for 15 min at 4°C). The supernatant was used to measure the VFAs. The VFA concentrations were determined by a capillary column gas chromatograph (13, 14).

For the enzyme activity assay, frozen pellets were thawed at room temperature. After being centrifuged at 3000g for 10 min (4°C), 10∼15 ml of supernatant was taken for sonication (power 400W, crushed 3 times, 30S each time, 30S interval), and the crushed liquid was the sample to be tested. The assayed CMCase, β-glucosidase, and xylanase activity measured used the 3,5-dinitrosalicylic acid method (15, 16).

The glucose, cellobiose and xylose content in the samples were determined with high-performance liquid chromatography (HPLC, Waters 600, USA) using the Aminex HPX-87H column (Bio-Rad, America) and a refractive index detector (Waters 2414, USA) with 0.005 M H_2_SO_4_ as the mobile phase, a column temperature of 60°C, and a velocity of 1.0 mL min^-1^, as assessed by a refractive index detector. The cellulose, hemicellulose, lignin, neutral detergent fiber (NDF) and acid detergent fiber (ADF) content were analyzed using the Ankom A200 fiber analyzer (Ankom Technology, Macedon, NY) using the method of Van Soest et al.(1991) (17). Briefly, the hemicellulose content was estimated as the difference between the NDF and the ADF, the cellulose content was estimated as the difference between the ADF and the acid-detergent lignin (ADL), and the lignin content was estimated as the difference between the ADL and the ash content.

### 2.3 DNA and RNA extraction

#### 2.3.1 DNA extraction

Genomic DNA was extracted according to An et al.(2005)(18)and Minas et al.(2011) (19)with some improvements. DNA extraction was performed based on a CTAB-based DNA extraction method. The CTAB lysis buffer contained 2% w/v CTAB (Sigma-Aldrich, Poole, UK), 100 mM Tris–HCl (pH = 8.0; Fisher), 20 mM EDTA (pH = 8.0; Fisher) and 1.4 M NaCl (Fisher). The pH of the lysis buffer was adjusted to 5.0 prior to sterilization by autoclaving (20). The final DNA was resuspended in 100 μL TE buffer (pH = 8.0; Sigma-Aldrich) and stored at −80°C.

#### 2.3.2 RNA extraction, RNA Reverse Transcribed and qPCR primer design and analysis

RNA extraction was performed used the liquid nitrogen grinding + TRIzol reagent (Ambion, Carlsbad, USA); the steps are as described by Kang et al.(2009) and Wang et al.(2011) (21, 22) with some improvements. The RNA was reverse transcribed cDNA using a PrimeScript™ 1st strand cDNA Synthesis Kit (Code No. 6110A, TAKARA, Dalian, China), following the kit instructions. The reverse transcribed PCRs were as follows: 37°C for 15 min, 85°C for 5 sec, and 4°C for 10 min. cDNA stored the rest at −80°C. The PCR primers were listed in Tab.1 and were assembled from the literature (23, 24). Primers were provided by Sangon Biotech (Shanghai) Co.,Ltd (Shanghai, China).

The number of rumen microorganisms is expressed as a percentage relative to the total rumen 16Sr DNA: target bacteria (% total bacterial 16Sr DNA) = 2^-(Ct^ ^target^ ^bacteria-Ct total bacteria)^ × 100%, where target is the specific microbial group of interest.

### 2.4 Deep Sequencing and KEGG Analysis

Illumina TruSeq libraries were prepared from genomic DNA and sequenced on an Illumina HiSeq 2500 instrument by Edinburgh Genomics. Five hundred bp paired-end reads were generated, resulting in between 8.08 and 10.09 Gb per sample (between 65.84 and 83.68 million paired reads). For a functional analysis, classification functions were classified using the KEGG orthology database (version 67.1) to identify relationships between various pathways and obtain KEGG numbers and EC numbers. First, we matched the reads directly to KEGG genes, and the mismatch is allowed to be within 10%. All KEGG Orthologue groups (KO) with a hit equal to the best hit were examined. If we were unable to resolve the read to a single KO, the read is ignored; otherwise, the read was assigned to a unique KO. A statistical analysis was performed on each microorganism or function using the PROC GLM of SAS 9.4.

### 2.6 Statistical Data Analyses

Cellulose, hemicellulose and lignin were analyzed used covariance, and the content of each component in the diets was recorded as a covariate. The ANOVA statistical analyses of VFAs, pH, lactate, enzyme activity, sugar content and microbial diversity were done using the PROC GLM of SAS 9.4. The treatment was considered a fixed effect. Statistical significance was declared at *P*<0.05.

## 3. RESULTS

### 3.1 Rumen fermentation parameters affected by different F:C ratio in diets

An overview of the analyses of in vivo fermentation, rumen pH, NH3-N, the content of VFA and lactate is provided in Tab.2. Rumen pH, NH3-N, the content of isobutyrate and lactate, and the ratio of acetate:propionate(A:P) showed a significant increase with the dietary forage level increase, while the content of TVFA, propionate, butyrate, valerate, and isovalerate significantly decreased (*P* < 0.01). Acetate was the only fermentation product that did not differ significantly between the HF group and HC group (*P* = 0.59). The content of TVFA in the rumen of the HC group was higher than that in the HF group, and the accumulation of VFA in the rumen resulted in a decrease in rumen pH. The concentration of NH_3_-N decreased significantly with the increase of concentrate in the diet.

### 3.2 The composition of diets and rumen contents

A covariance analysis was used to analyze the composition of rumen content in the HF and HC groups. The chemical composition of rumen contents was shown in Tab.3. The cellulose, hemicellulose, and acid detergent lignin (ADL) in the diets of HF group were significantly higher than in the HC group, but in the rumen content, there was no difference between two groups. In the rumen content, the HC group has a higher (*P* < 0.05) content of cellobiose (Fig.2A) and glucose (Fig.2B) compared to the HF group. The higher content of cellobiose and glucose indicate that the releasing rates of these sugars were faster from the HC group than from the HF group. This phenomenon is consistent with the β-glucosidase (Fig.2D), and the β-glucosidase activity in the HC group was higher than the HF group. The fiber digestibility of HC group was shown to be higher than of HF group, because the content of cellulose in the HC group was lower, and the content of the cellobiose decomposed by cellulose and glucose decomposed by cellobiose were higher than the HF group. Moreover, the carboxymethyl cellulose (CMCase) (Fig.2C) and β-glucosidase activity in the HC group were higher than in the HF group. The content of xylan and xylose were higher in HF group but not significantly so (Fig.3A,3B). The β-xylosidase in the HC group was significantly higher than in the HF group (Fig.3C).

**Fig. 2.**
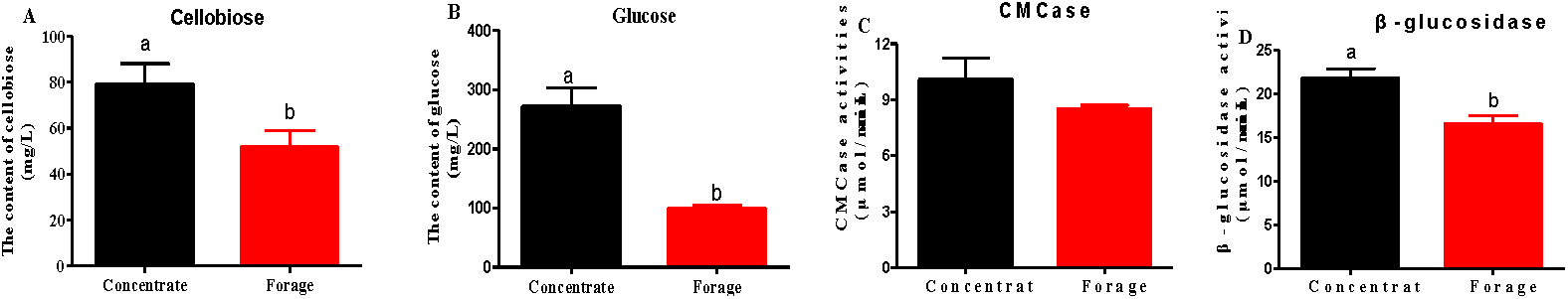
Cellulose decomposition of rumen contents. (A) Cellobiose, (B) Glucose, (C) CMCase, (D) β-glucosidase. The error bars represent the standard error of the mean (n = 3). Different letters in each figure panel indicate a significant difference (P < 0.05).

**Fig. 3.**
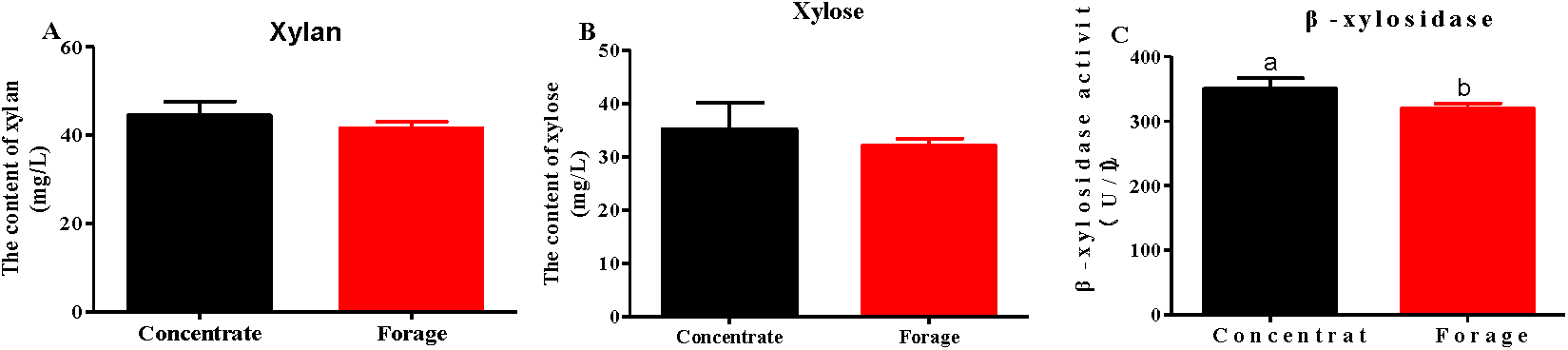
Hemicellulose decomposition of rumen content. (A) Xylan, (B) Xylose, (C) β-xylosidase. The error bars represent a standard error of the mean (n = 3). Different letters in each figure panel indicate a significant difference (P < 0.05).

### 3.3 Ruminal microbial community changes within HC and HF groups

At a threshold of >0.1% relative abundance, 57 bacterial taxa (97% sequence similarity) were retrieved from the 16S rDNA gene amplicon sequences, while 52 taxa were retrieved from the metagenome genes. At the family level, the five taxa accounted for approximately 65% of the 16S rRNA gene amplicon sequences and the metagenome data in all samples: *Acidaminococcaceae, Lachnospiraceae, Prevotellaceae, Ruminococcaceae* and *Succinivibrionaceae* (Fig.4). Although there was no significant difference (P ≤0.05) in the abundance of *Ruminococcaceae* between HF and HC groups. All other taxa showing different abundance at the family level, *Lachnospiraceae* and *Succinivibrionaceae* being more abundant in the HF group, yet, *Prevotellaceae* and *Acidaminococcaceae* being more abundant in the HC group, that all based on both 16S rDNA amplicon and metagenomic sequencing data (Fig. 4a and 4b). At the genus-level resolution, eight taxa presented significantly different relative abundances in the HF and HC groups based on GLM (P ≤ 0.05), four of which were more abundant in the HF group and four were more abundant in the HC group (Tab.4). The most noteworthy were *Prevotella* (family *Prevotellaceae*), *Selenomonas_1* and *Veillonella* (both from the family *Acidaminococcaceae*), which were more abundant in the HF group based on both 16S rDNA and metagenome datasets and occupy 3.47, 2.42 and 1.43% of the metagenome reads from the HC group, respectively. *Ruminococcus* (family *Lachnospiraceae*) were the principal rumen cellulose-degrading bacteria and were significantly more abundant in the HF group in both datasets, with an average relative abundance of ∼2% in these animals. *Prevotella* and *Selenomonas_1* belong to the principal rumen starch-degrading bacteria. *Succinivibrio* and *Selenomonas* in the HC group were more enrichment may be related to the diet in HC group has more starch and maltose and less cellulose and xylan.

**Fig. 4.**
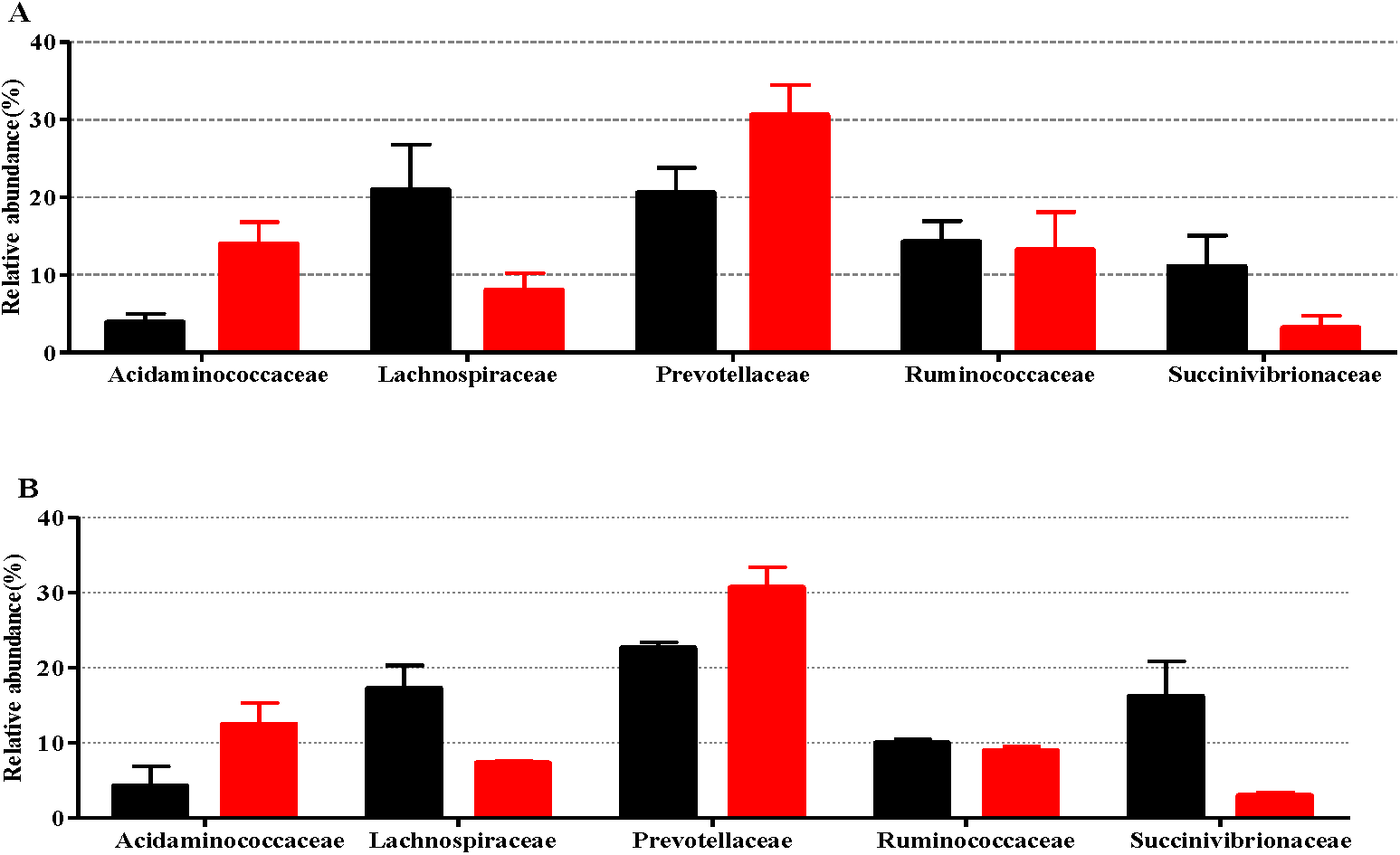
Relative abundance of the most highly represented bacterial families based on 16S rDNA gene amplicon sequencing data (a) and 16S rRNA genes retrieved from the metagenome dataset (b) from rumen content samples of the HF group (black) and HC group (red). **P < 0.01, *P < 0.05. Error bars denote one standard deviation.

### 3.4 Volatile fatty acid metabolism

Volatile fatty acids act as energy sources for ruminants; therefore, it was important to study the metabolism of VFAs. The production of acetate was contained in the metabolic pathway of butyrate; therefore, the metabolism of propionate and butyrate was the main research focus.

#### 3.4.1 Genes directly involved in propionate

Many genes involved in the metabolism of short chain fatty acids are diverse and abundant in the metagenomic dataset, including genes encoding pyruvate fermentation to lactate and further fermentation to propionate, which transformation pyruvate to propionate, including the acrylate pathway and the succinate pathway. Plant polysaccharides are fermented by the rumen microbial, and finally, propionate is mainly production.

Genes encoding enzymes that are directly involved in propionate were analyzed for their abundance in HC and HF groups. Genes encoding the lactate-acrylate pathway produce propionate (Fig.S1). With the exception of the gene K01026 for lactyl transferase (EC:2.8.3.1), lower abundance in the HF group was found. The other genes, K00016 for acyl-CoA dehydratase (EC:1.3.8.7) and K00249 for lactate dehydrogenase (EC:1.1.1.27), directly involved in propionate were higher in the HC group than in the HF group. However, in the lactate-acrylate pathway, the representative genes of lactoyl-CoA dehydratase (EC: 4.2.1.54) are not included in the KEGG gene dataset; therefore, this metabolic pathway cannot produce propionate. Conversely, fermenting to propionate seems to be more likely given the high readings of the genes involved via the succinate pathway. Such a pathway has been demonstrated in Fig.5. Genes involved in the succinate pathway to propionate showed significantly higher read counts in the HC group, which included genes K01902 and K01903 for succinate-CoA synthetase (EC:6.2.1.5) and gene K01026 for lactyl transferase (EC:2.8.3.1), both of which were predicted as being the limiting factors for propionate formation. Pyruvate carboxylase (EC 6.4.1.1) of genes K01958, K01959 and K01960 was higher in the HF group than in the HC group. The other intermediate enzymes were all found, and the difference was not significant. VFA produced by rumen fermentation can be used as an energy supply for animals, of which only propionate gluconeogenesis is the main source of glucose. Therefore, propionate has important physiological significance for ruminants.

**Fig. 5.**
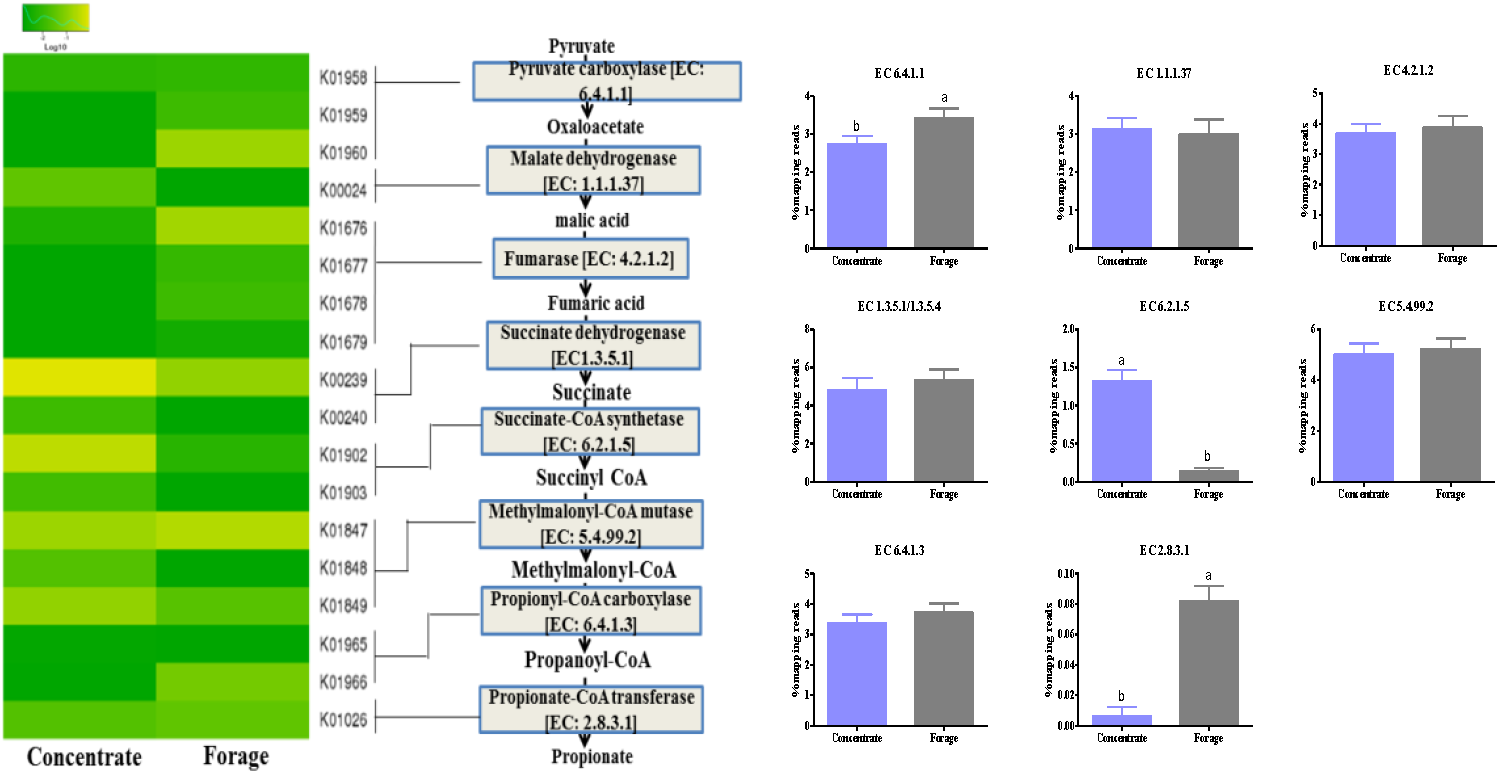
The metagenomic abundance of key elements of the propionate production pathway. Center pane: the propionate production pathway showing enzyme classification (EC) numbers. Left pane: heat map of KEGG orthologues for the EC numbers involved in propionate production (lines connect the heat map to the propionate production pathway indicating which K0 numbers represent the given enzymes). Right pane: the abundance of each of the relevant EC numbers in our data set. The bar charts show the percentage of reads mapped to each enzyme in the 2 groups (HC and HF groups). The blue bars are cattle selected for HC groups, and gray bars are cattle selected for HF groups.

#### 3.4.2 Genes directly involved in butyrate

The genes encoding the butyrate formation pathway were analyzed for their abundance in the HC and HF groups (Fig.6). With the exception of gene K00248 for butyryl-CoA dehydrogenase (EC:1.3.8.1) and genes K01034 and K01035 for acetate-CoA transferase (EC:2.8.3.8), the content of all genes encoding the translation of pyruvate into butyrate in HC group showed higher or not significantly different than in HF group, at gene levels. The genes for acetoacetyl-CoA reductase (EC:1.1.1.36) and correlations to butyrate yield were not available in the KEGG database. The enzymes of butyryl-CoA dehydrogenase (EC:1.3.8.1) and acetate-CoA transferase (EC:2.8.3.8) were needed for the last two steps in the formation of butyrate, Therefore, the butyrate in the HF group should be higher than that in the HC group, which was consistent with the content of butyrate determination. No difference was found in Phosphate butyryltransferase (EC:2.3.1.19) between the HF and HC groups. Gene K00929 for butyrate kinase (EC:2.7.2.7) in the HC group was higher than HF group. From fig.6, we can see that butyrate kinase (EC:2.7.2.7) is the finale enzyme to produce butyrate from another branch pathway. Therefore, butyryl-CoA directly produces butyrate via acetate-CoA transferase (EC: 2.8.3.8), rather than another branch pathway. This result may be due to the high content of rumen microbes that produce butyrate or use of acetate as a precursor for butyrate synthesis under conditions of lactate fermentation (where lactate fermentation could occur).

**Fig. 6.**
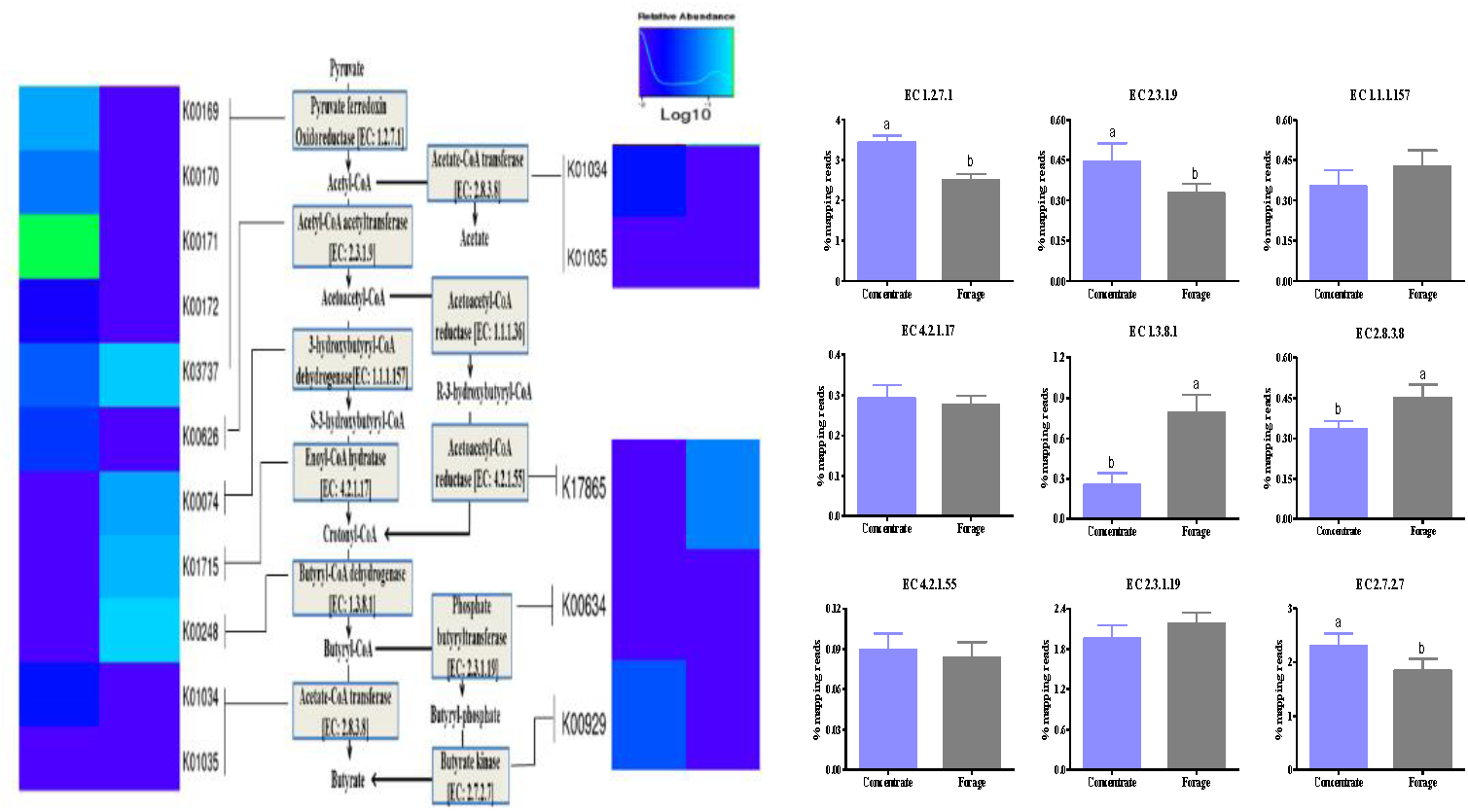
The metagenomic abundance of key elements of the butyrate production pathway. Centre pane: the butyrate production pathway, plus ancillary reactions, showing enzyme classification (EC) numbers. Left and right pane: heat map of KEGG orthologues for the EC numbers involved in butyrate production (lines connect the heat map to the butyrate production pathway, indicating which K0 numbers represent the given enzymes). Lower pane: the abundance of each of the relevant EC numbers in our data set. Bar charts show the percentage of reads mapped to each enzyme in the 2 groups for diet (high or low concentrate). Grey bars are cattle selected for HF groups and blue bars are cattle selected for HC groups.

### 3.5 The major microbial material involved in cellulose and volatile fatty acid metabolism gene characterization and quantification

The amount of major microbial material involved in cellulose and VFA metabolism genes was estimated from the relative abundance of rumen total bacteria, which was determined by qPCR using specific bacterial primers of HF and HC groups. In our study, we selected eight representative genes (Tab. 5). At present, *Ruminococcus albus* and *R. flavefaciens* are considered to be the major cellulolytic bacteria in the rumen (25), and the relative abundance in the HF group is higher than in the HC group. *Ruminococcus albus* and R. *flavefaciens* both have relative content at approximately 7%. *Fibrobacter succinogenes* and *Butyrivibrio fibrisolvens* are the main cellulolytic bacterial, and the relative content of the HF group was significantly higher (*P*≤0.05) compared to that of the HC group. These results confirmed the previous research in this study on cellulose and enzyme activity. The relative content of the species *Prevotella ruminicola, Selenomonas ruminantium*, and *Megasphaera elsdenii* were higher in the HC group than in the HF group.

**Table 5.** Differences in the relative expression (%) of the main bacteria in the HF and HC groups. The relative expression (%) levels are shown.

| Main microbial | Forage group | Concentrate group | SEM | *p*-value |
| --- | --- | --- | --- | --- |
| *Ruminococcus flavefaciens* | 5.0003 | 3.8988 | 0.4994 | 0.1499 |
| *Ruminococcus albus* | 2.2981 | 2.7131 | 0.4303 | 0.5207 |
| *Fibrobacter succinogenes* | 0.4214 | 0.3268 | 0.0691 | 0.0032 |
| *Butyrivibrio fibrisolvens* | 0.0486 | 0.0201 | 0.0065 | 0.0112 |
| *Prevotella ruminicola* | 2.7294 | 4.3986 | 0.3606 | 0.0113 |
| *Selenomonas ruminantium* | 1.6891 | 3.7616 | 0.2054 | 0.0022 |
| *Megasphaera elsdenii* | 0.0191 | 0.0034 | 0.0008 | 0.0001 |
| *Veillonella alkalescens* | 0.0339 | 0.2068 | 0.0320 | 0.0088 |

## 4. DISCUSSION

### Rumen fermentation parameters and rumen contents

In the present study, we investigated the fermentation parameters, Our findings show that HC diet produce more VFAs (Tab.2), and the accumulation of VFAs in the rumen resulted in a decrease in rumen pH (26). The concentration of NH3-N decreased significantly with the increase of concentrate in the diet. Michalski et al.(2014)(27) found that non-fibrous carbohydrates (monosaccharides, disaccharides, polysaccharides, etc.) in the diet are the main factors limiting the utilization of NH3-N by rumen microbes and that raising the level of nonfibrous carbohydrates in the diet can promote utilization of NH3-N by rumen microbes.

In the rumen content, the HC group has a higher content of cellobiose and glucose compared to the HF group. Cellobiose generates glucose by β-glucosidase, Chen(2012) (28) showed that β-glucosidase is a key factor in the conversion of cellobiose to glucose and enhancing the efficiency of cellulolytic enzymes for glucose production. In this study, the cellulose digestibility of HC group was shown to be higher than for HF group may be due to the content of cellulose in the HC group was lower and the content of the cellobiose decomposed by cellulose and glucose decomposed by cellobiose were higher than the HF group.

Xylan is the main component of hemicellulose; therefore, the degradation of hemicellulose was studied with xylan changes. In our research, the β-xylosidase in the HC group was significantly higher than in the HF group. β-xylosidase is an exo-enzyme that mainly catalyzes the hydrolysis of xyloside and from the nonreducing hydrolysis xylo-oligosaccharide to xylose (29, 30), which can effectively relieve feedback inhibitions of xylanase by endo-xylanase hydrolysates (xylo-oligosaccharides) (31). Hemicellulose digestibility of HC group was shown to be higher than for HF group, which is consistent with the digestibility of cellulose.

### 4.2 Ruminal microbial community changes within HC and HF groups

At the family level, The rumen microbial community comprises mainly *Acidaminococcaceae, Lachnospiraceae, Prevotellaceae, Ruminococcaceae* and *Succinivibrionaceae*, which accounted for approximately 65%. At the genus level, our study focused on the main and different microbial other than the all microbial. *Prevotella* (family *Prevotellaceae*), *Selenomonas_1* and *Veillonella* (both from the family *Acidaminococcaceae*). *Prevotella*, as part of the core microbiome, can grow rapidly on starch media and produce final products other than lactate (mainly succinate and propionate) (32). The reason for the enrichment of *Succinivibrio* and *Selenomonas* in the HC group may be related to the diet in which the HC group has more starch and maltose and less cellulose and xylan. Previous researches (33-35) have shown that *Succinivibrio* and *Selenomonas* growth on starch, maltose and soluble sugars and cellulose and xylan are not available, which produces more succinate, which decarboxylates and leads to more propionate formation (36). Recently, extensive investigations have been carried out on the microbial communities of multiple groups of cows with different F:C ratios. *Ruminococcus* (family *Lachnospiraceae*) were the principal rumen cellulose-degrading bacteria. Cerrillo et al. (1999) (37) reported that the forage group was abundant in *Ruminococcus*, among other cellulose-degradation bacteria, and studies have shown that *Ruminococcus* are fermented with a large amount of cellulose as a substrate and can form more acetate (38, 39), which is consistent with the results of our study of the forage group to produce more acetate and have more cellulose and hemicellulose.

### 4.3 Volatile fatty acid metabolism

Volatile fatty acids act as energy sources for ruminants; therefore, it was important to study the metabolism of VFAs. The production of acetate was contained in the metabolic pathway of butyrate; therefore, the metabolism of propionate and butyrate was the main research focus.

#### 4.3.1 Genes directly involved in propionate

In rumen, plant polysaccharides are fermented by the rumen microbial, propionate is one of the final production. Propionate can be produced by acrylate pathway and succinate pathway. The acrylate pathway is the primary pathway in the case of animal diets with high starch content (40, 41), and *Megasphaera elsdenii* is the major propionate producer via the acrylate pathway (42, 43), in our study, the content of *Megasphaera elsdenii* was very low at the expression (Tab.5) and genes for lactyl-CoA dehydratase was not found. Conversely, fermenting to propionate seems to be more likely given the high readings of the genes involved via the succinate pathway. The CO2 was immobilized in phosphoenolpyruvate to form oxaloacetate, which was then succinate produced by malate and fumarate, and succinate is rapidly converted to propionate by the microbial enzyme (32). Such a pathway has been demonstrated in *Veillonella* (44) (Fig. 4). Studies have shown that *Bacteroides ruminicola* with glucose is the main fermentation substrate, and the main fermentation products are succinate, CO2, formate and acetate (45, 46). In vitro fermentation studies have shown that succinate as a result of carbohydrate fermentation is rapidly converted to propionate in the rumen (32). *Selenomonas ruminantium*, another rumen species, can also produce propionate by fermenting carbohydrates or lactate, or produce propionate by decarboxylation of succinate. VFA produced by rumen fermentation can be used as an energy supply for animals, of which only propionate gluconeogenesis is the main source of glucose. Therefore, propionate has important physiological significance for ruminants.

#### 4.3.2 Genes directly involved in butyrate

The content of butyrate in HF group was higher than HC group, this result was consistent with Mccullough and Sisk research. Mccullough and Sisk (47) studied the effects of different diets on rumen VFAs and found that the butyrate yield in the forage group was higher than that in the grain group. This result may be due to the high relative content of rumen microbes that produce butyrate. Butyrate production and accumulation appear to increase when high-fiber degradability coincides with the high availability of nonstructural carbohydrates (48). This also may be related to the use of acetate as a precursor for butyrate synthesis under conditions of lactate fermentation (where lactate fermentation could occur). Under these conditions, lactate-producing bacteria such as *Butyrivibrio fibrisolvens* can directly produce butyrate using acetate via butyryl-CoA/acetic acid-CoA transferase (EC: 2.8.3.8) rather than by converting two acetyl-CoA molecules into acetoacetyl-CoA (49). The *Butyrivibrio fibrisolvens* relative content at the transcriptional level from our study was similar to the findings of Diezgonzalez et al. (1999), which was the HF group with the higher *Butyrivibrio fibrisolvens* relative content, and then the butyrate of the forage group was higher. On the other hand, *M. elsdenii* produces butyrate via the malonyl-CoA pathway from various reactions involving acetyl-CoA, which is activated by acetate and is combined with CO_2_ to form malonyl-CoA (50). The lactate fermentation of *M. elsdenii* is not regulated by glucose or maltose, so the utilization of lactic acid increases with the feeding of soluble sugar (43, 51). Others have also reported that when *M. elsdenii* is purely cultured, the accumulation of butyrate prevented high levels of lactate accumulation. The magnitude of the effect was positively correlated with the dose of *M. elsdenii*. Our recent work indicated that the relative content of *M. elsdenii* was very low in both HF and HC groups; however, under this condition, fermentation to butyrate seems more likely via the butyryl-CoA/acetate-CoA transferase pathway using acetate as an acceptor.

### 4.4 The major microbial material involved in cellulose and volatile fatty acid metabolism gene characterization and quantification

Abundance of microbial involved in cellulose and VFA metabolism encoding genes was measured by quantifying the microbial gene copy number as well as the expression level. Based on the DNA level, different patterns of abundance of microbial encoding genes were found in HF and HC groups. *Ruminococcus albus* and R. *flavefaciens* both have relative content at approximately 7%. Previous studies (17, 25) have shown that they constitute up to approximately 10% of the total bacterial isolates in either HF or HC diets, and increasing rumen *Ruminococcus* increases the ratio of A:P. Our present study was consistent with Weimer(1998) (52).

The enzyme produced by the *Fibrobacter succinogenes* was confirmed to have an independent cellulose catalytic zone and a cellulose binding zone (53) and has a strong ability to degrade plant cell walls. The *Fibrobacter succinogenes* can produces a variety of β-glucanases, which degrade cellulose and xylanase, which in turn degrades hemicellulose. *Butyrivibrio fibrisolvens*, another rumen species, produces xylanase and endoglucanase (54). These results confirmed the previous research in this study on cellulose and enzyme activity. *Prevotella ruminicola* is a dominant bacterium in the rumen and specifically degrades the noncellulosic components in the plant cell wall and can simultaneously metabolize pentose and glucose but first utilizes pentose and then cellobiose (55). Therefore, in the rumen, we presumed that the pentose (main xylose) content is lower than the cellobiose content. Our observations confirm that the content of xylose (Fig.2) was lower than cellobiose in the rumen (Fig.1). *Selenomonas ruminantium* is unable degrade cellulose directly, but it can produce succinate by using the cellobiose of the cellulose degradation product, and *Selenomonas ruminantium* and *Veillonella alkalescens* via the succinate pathway produce propionate (56). *Selenomonas ruminantium* can also produce acetate via the acetyl-CoA pathway, so it plays an important role in the metabolism of propionate. In the rumen, *Megasphaera elsdenii* is regarded as the main fermenter of lactate (51) under the conditions of rapid fermentation of sugar and production of more lactate, which results in the relative content in the bacterial community increasing (57). In our present study, the relative content of *Megasphaera elsdenii* in the HFgroup was higher than that in the HC group, and coincident with that, the lactate content was also higher in the HF group, However, the HC group should contain more sugar, and the relative content of *Megasphaera elsdenii* was higher, which may be related to our sampling time and the interaction between microorganisms, which requires further research.

## 5. CONCLUSION

In conclusion, this study combined metagenomics and metabolism to explore the effects of HF and HC diets on the cellulose degradation process and ruminal microbial communities in rumen. Feeding a HF diet increased ruminal pH and decreased TVFA concentration. The content of *Ruminococcus, Papillibacter* and *Roseburia* in HF group were higher, which could efficiently degrade cellulose in rumen, thereby enhancing the activity of CMCase by promoting microorganisms producing this enzyme. We discover that the butyryl-CoA dehydrogenase (EC:1.3.8.1) is the restriction enzyme in butyrate metabolism and succinate-CoA synthetase (EC:6.2.1.5) and lactyl transferase (EC:2.8.3.1) are the key enzyme in pyruvate metabolism, can reveal how fiber degradation and VFAs production are manipulated by metabolic pathways and microbial communities. Therefore, combined with cellulose, enzymes and VFA measurements, rumen microbiome and fermentate characterization will be a useful screening tool for choosing cellulolytic bacteria.

## ACKNOWLEDGEMENTS

The authors thank Long Jin at the Research and Development Center for their technical assistance. Hongjian Xu providing language help.

